# Persistence phenotype of adherent-invasive *Escherichia coli* in response to ciprofloxacin, revealing high-persistence strains

**DOI:** 10.1101/2024.11.05.622023

**Authors:** Valeria Pérez-Villalobos, Roberto Vidal, Marcela A. Hermoso, Paula Bustamante

## Abstract

Persister cells correspond to a bacteria subpopulation able to survive antibiotic treatments and are thought to contribute to chronicity and symptom relapse of chronic diseases. Crohn’s disease (CD) is a multifactorial chronic inflammatory condition of the gastrointestinal tract, and adherent-invasive *Escherichia coli* (AIEC) has emerged as a key contributor to its pathogenesis. AIEC can survive, replicate, and produce persister cells within macrophages; however, out of the LF82 reference strain, a lack of knowledge of the persistence phenotype and its variability among AIEC strains is lacking. Here, two AIEC reference strains were analyzed after ciprofloxacin challenge, a fluoroquinolone antibiotic common in CD treatments. Four clinical isolates with an AIEC phenotype and five different *E. coli* pathotypes were also analyzed. We investigated the roles of the resident antibiotic resistance plasmid and the stress response protein HtrA, along with macrophage-induced persister formation. We observed a broad variability of persister cell formation capability among AIEC strains. Remarkably, the reference NRG857c strain showed a threatening high-persistence phenotype, with persistence levels higher than LF82 (65-fold) and some AIEC clinical isolates. The resident antibiotic resistance plasmid and the stress protein HtrA were dispensable for the NRG857c persistence phenotype. Among *E. coli* pathotypes, only EPEC showed persistence levels comparable to NRG857c, while other pathotypes resembled the lower persistence levels of LF82. Macrophage internalization did not increase NRG857c persister fraction, opposite to the LF82 behavior. Overall, our findings show that variable persistence phenotypes are found among AIEC. Opposite to LF82, NRG857c is a high-persistence strain, and this characteristic is shared with one AIEC clinical isolated and the EPEC reference strain. Undoubtedly, bacterial persistence should be considered in CD antibiotic treatments, especially with the feasible presence of AIEC high-persistence strains.

**Author Summary:** Adherent-invasive *Escherichia coli* (AIEC) are common bacteria found in Crohn’s disease patients and recognized to survive within macrophages. The reference strain LF82 is known to form persistent cells, an enigmatic subpopulation of cells able to survive antibiotic treatments and thought to contribute to chronicity and symptoms relapses of chronic diseases. However, out of LF82, there is a lack of knowledge on AIEC persister cell formation, as well as the genetic factors contributing to this phenotype. In this study, it was observed a that the persistence phenotype was variable among both AIEC strains and *E. coli* pathotypes. Unlike LF82, the reference strain NRG857c displayed inherently high-persistence levels, regardless of whether it carries its native antibiotic resistance plasmid or the *htrA* and *hipA* genes. Notably, stress conditions within macrophages did not trigger increased persister cell formation in NRG857c. This study highlights the importance to decipher the role of AIEC persisters in ongoing Crohn’s disease, symptom relapses, and responses to antibiotic treatment, especially given the potential existence of high-persistence AIEC strains.

## Introduction

Recurrence of symptoms in chronic diseases is mainly due to relapse rather than reinfection [1], and a subpopulation of transient antibiotic tolerant bacteria, known as persister cells, are thought to be key actors in these processes [1–3]. Persister are slow-growing or growth-arrested bacterial cells, with a decreased but still active metabolism [4], whose formation has been linked to the stringent response through (p)ppGpp, the SOS response, and toxin-antitoxin systems [2,5], without lack of controversy [6,7]. In addition, mutations can increase the level of persistence, as the recognized *hipA7* variant [8] and other high-persistence mutants observed in patients subjected to repeated antibiotic treatments [9,10].

The presence of persisters during infections has been observed in adherent-invasive *E. coli* (AIEC) [11], a diverse *E. coli* pathotype with high prevalence in Crohńs disease (CD) patients [12,13]. Bacterial contribution is key for the onset of CD, promoting chronic inflammatory relapses [14], and consequently, ciprofloxacin and/or metronidazole treatments have shown positive results in clinical trials [15].

Pathogenic mechanisms of AIEC are elusive, but it is characterized by its ability to adhere and to invade intestinal epithelial cells, and colonize macrophages [16,17]. Reference AIEC strains, LF82 and NRG857c, have evolutionary relationship [18], and strain-specific genetic elements encoded on the chromosome and on large extrachromosomal plasmids unique to each isolate. A main characteristic of the pathotype is the ability to survive and replicate within macrophages [17], where the protease HtrA plays an important role [19]. Besides, macrophages induce the formation of LF82 persister cells [11].

Considering the diversity found among AIEC members and that persistence has been studied exclusively in the LF82 strain, the aim of this study was to analyze the persistence phenotype of AIEC reference strains and clinical isolates, comparing with other *E. coli* pathotypes, and to assess whether chromosomal or extrachromosomal genetic factors were involved, along with the effect of macrophage in formation of AIEC persister cells.

## Results

### Reference AIEC strains have variable persistence phenotypes

We analyzed the persistence phenotype of reference AIEC strains (S1 Table), in response to ciprofloxacin, an antibiotic commonly used in CD treatment [15]. Although both strains behaved with the characteristic biphasic killing curve of persister cell formation, their surviving fractions after the ciprofloxacin challenge were significantly different (Fig 1A). Remarkable, at 5-hour post-antibiotic challenge ∼0.13% of the NRG857c population survived, 65-fold compared to LF82 at the same time point. Of note, we got similar survival fractions for NRG857c when a MOPS-minimal medium was used, even with lower antibiotic concentrations (S1 Fig). Surviving NRG857c bacteria did not acquire antibiotic resistance during the experiment (data not shown) and behaved as the original bacterial culture following antibiotic challenge (S2 Fig), revealing they are reliable persister cells. Overall, our results show that AIEC reference strains have contrasting persistence phenotypes, with NRG857c having a remarkable high persistence level.

**Figure 1.**
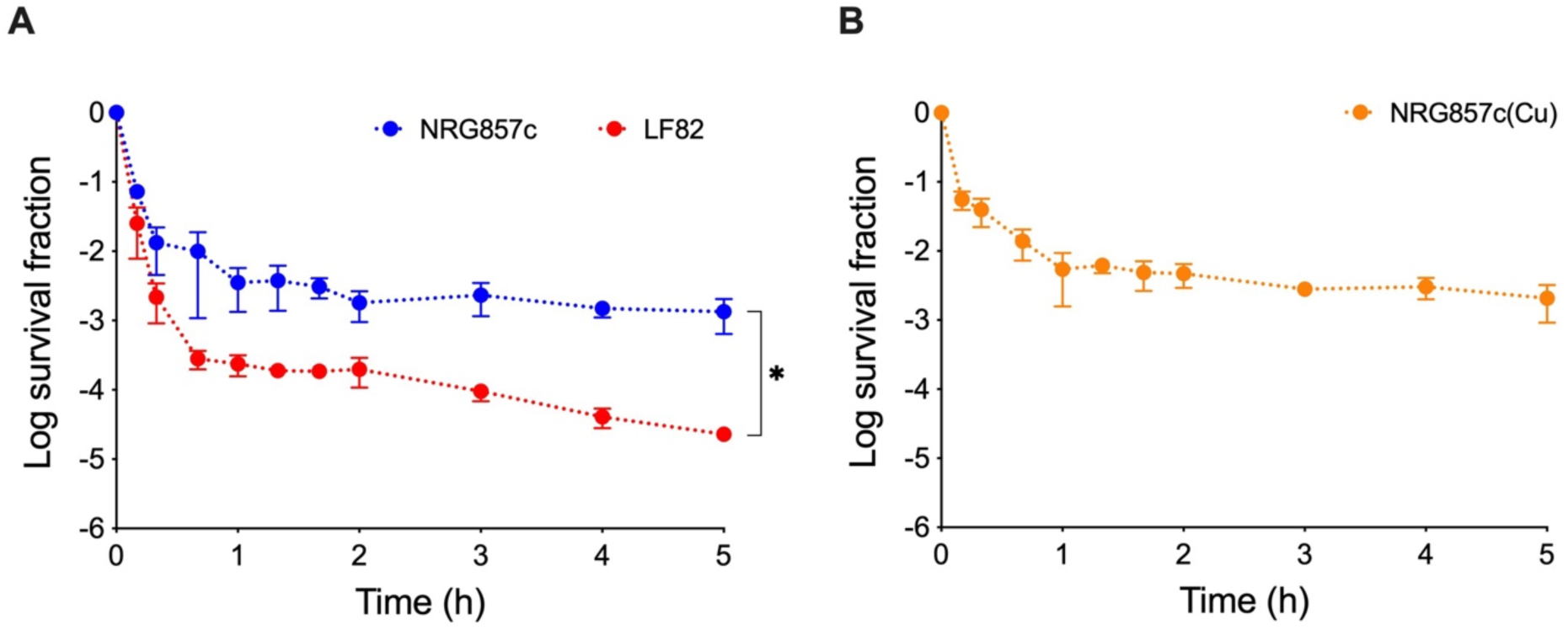
Time-killing curves of reference AIEC strains. (A) NRG857c and LF82 strains or (B) NRG857c(Cu) strain, a derivative of NRG857c cured of its multi-resistance plasmid, were grown in LB broth, challenged with 0.6 µg/ml ciprofloxacin, and survival was monitored at indicated times. Data points are mean values of three independent experiments, and error bars represent SDs. Student’s t-test was performed for NRG857c and LF82 strains data at 5-hours post treatment (*P < 0.05).

### The NRG857c multi-resistance plasmid is dispensable for persistence

NRG857c harbors a native multi-resistant plasmid, pO83_CORR [18], but its absent on a plasmid cured strain [20] did not affect growth (S3 Fig), neither ciprofloxacin MIC value (S2 Table). In addition, we found that the high-persistence phenotype of NRG857c does not rely on the carriage of pO83_CORR, or any gene encoded by it (Fig 1B), as ∼0.21% of the plasmid cured strain population survived after 5-hour post-antibiotic treatment, a level comparable to the wild-type strain.

### The AIEC pathotype shows variable persistence phenotypes

To elucidate if variability in persistence seen between AIEC reference strains extends to other AIEC members, we tested survival of four *E. coli* clinical isolates with an AIEC phenotype, CD1a, CD2a, CD6b and CD6r [21] (S1 Table) (Figs 2A-2B). Our results revealed that none of the AIEC clinical isolates reached persistence higher than NRG857c (Fig 2A). However, at 5-hour post-treatment, CD2a and CD6b isolates behaved without significant differences in their survival fractions in comparison to NRG857c (Fig 2B). On the contrary, CD1a and CD6r isolates resembled LF82 behavior (Fig 2B). These findings reveal that AIEC cover bacteria with variable persistence phenotypes, with the worrying occurrence of high-persistence strains.

**Figure 2.**
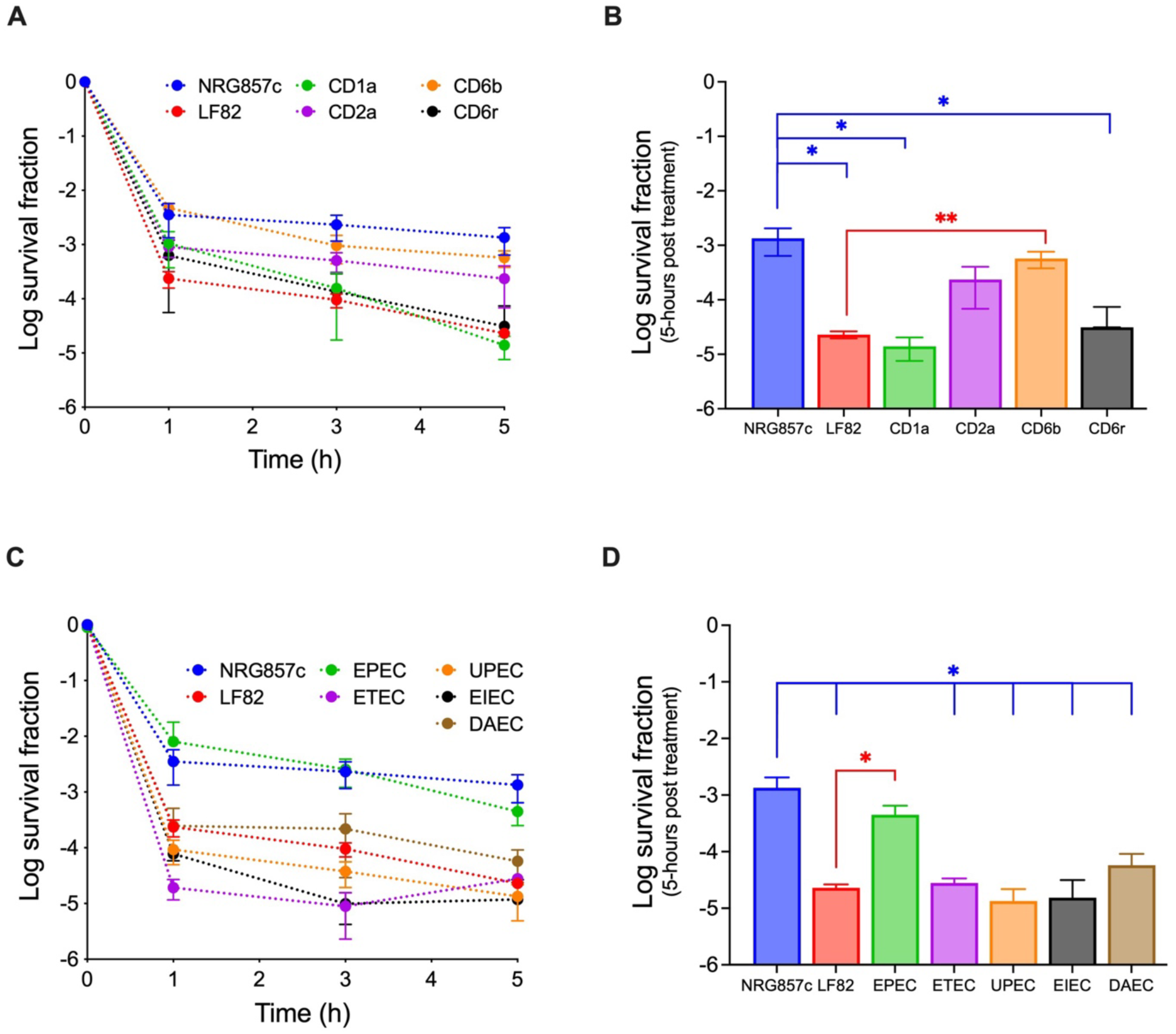
Time-killing curves of AIEC clinical isolates and *E. coli* pathotypes. (A, B) *E. coli* clinical isolates showing an AIEC phenotype, or (C, D) *E. coli* strains belonging to different pathotypes, were grown in LB broth, challenged with ciprofloxacin and survival was monitored at indicated times. The NRG857c and LF82 data from Fig. 1A, were included for reference (blue and red lines, respectively). (B, D) Details of survival data at 5-hours post-treatment for each strain graphed in A and C, respectively. Data points are mean values of three independent experiments, and error bars represent SDs. Student’s t-test was performed between NRG857c or LF82 and the other strains (* P < 0.05, ** P < 0.01).

### Persistence levels of NRG857c are the highest among *E. coli* pathotypes

To decipher if NRG857c persistence levels could also be achieved by other *E. coli* pathotypes, or they are a distinctive feature of some AIEC strains, killing curves of ETEC, EPEC, UPEC, EIEC and DAEC pathotype reference strains were analyzed (S1 Table and Fig 2C-2D). As before, almost none of the reference strains reached persistence higher than NRG857c. Only EPEC showed a level comparable to NRG857c (Fig 2C, green and blue lines, respectively) and significantly different to LF82 (Fig 2D). ETEC, UPEC, EIEC and DAEC, reached survival fractions comparable to LF82 and significantly different to NRG857c (Fig 2D). Altogether, our results show that persistence levels of NRG857c are the highest between different *E. coli* pathotypes.

### NRG857c persistence levels remain unaltered after macrophage passage

*Salmonella* internalization into macrophage is required to stimulate in vitro persister cell formation [4], and a similar behavior has been observed with LF82 [11]. Surprisingly, our findings revealed that macrophage internalization did not increase the formation of NRG857c persister cells (Fig 3A). As our results showed increased persistence levels when LF82 was internalized by macrophages (Fig 3B), in line with a previous report [11], we discarded an experimental error in our macrophage infection protocol. HtrA is a protease that plays an essential role in AIEC intra-macrophage existence [19]. However, deletion of *htrA* did not affect the persistence levels in both AIEC references strains (Fig 3C). Remarkable, even after macrophage passage, NRG857c Δ*htrA* did not show differences with the wild-type strain (Fig 3D). Altogether our results suggest that NRG857c seems to have a high basal persistence level and, unlike *Salmonella* or LF82, is unaltered by stress conditions found inside the macrophage. Indeed, despite its essential role in intramacrophage and stress survival, HtrA seems not be involve in the persistence phenotype.

**Figure 3.**
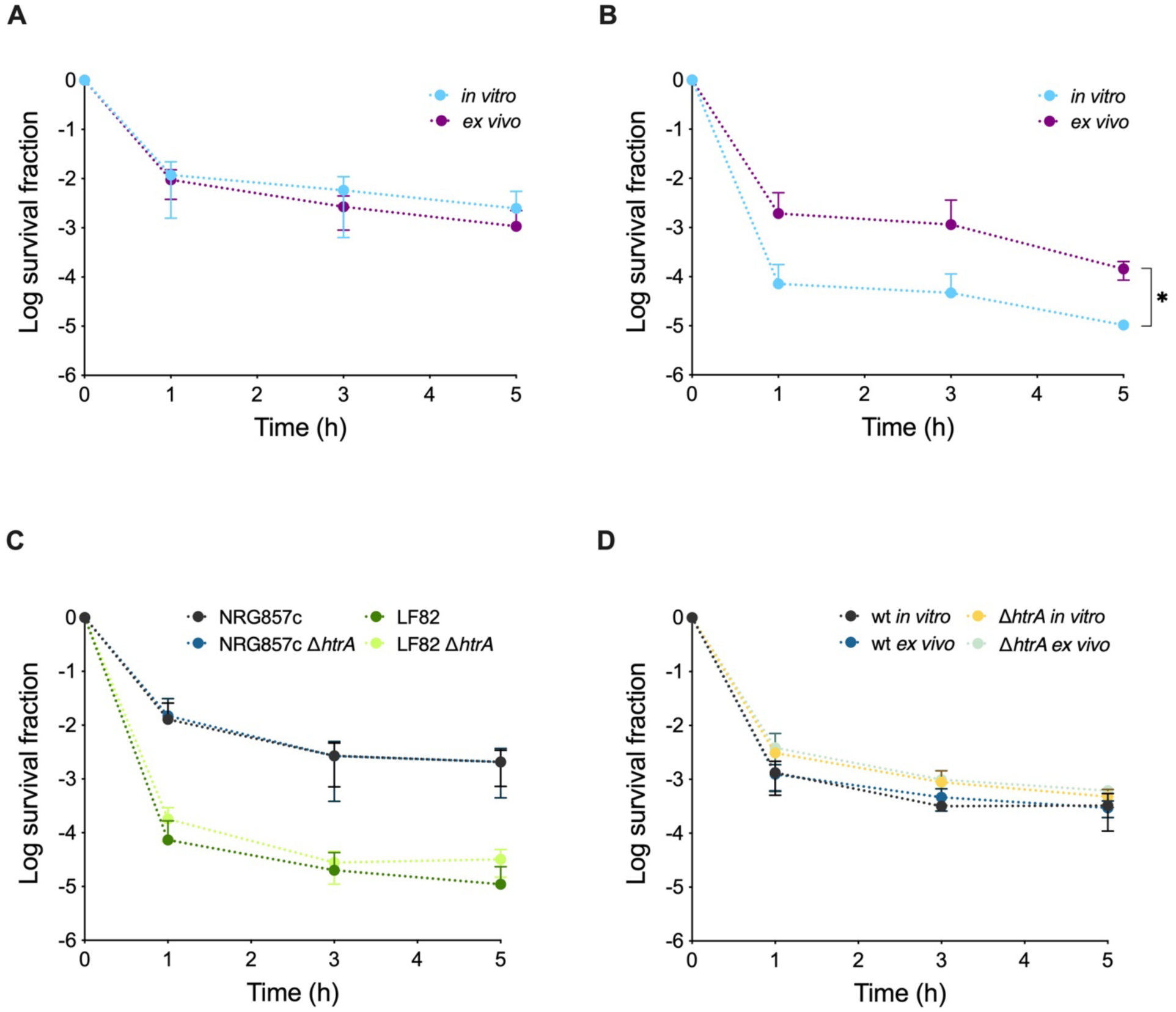
Killing curves of AIEC reference strains and their *htrA* deletion mutants after macrophage passage. (A) Wild type NRG857c and (B) LF82 were cultivated up to OD_600nm_ 0.3 in LB broth (in vitro, light blue lines) or harvested after 30 min post-infection within macrophages (ex vivo, purple lines), challenged with ciprofloxacin and survival was monitored at indicated time points. (C) In vitro killing curves of wild type reference AIEC strains and their *htrA* deletion mutants; the strains were grown in LB broth, challenged with ciprofloxacin, and survival was monitored at indicated times points. (D) Killing curves of wild type NRG857c and its *htrA* deletion mutant after macrophage passage. Same protocol as in A and B was followed. Data points are mean values of three independent experiments, and error bars represent SDs. Student’s t-tests was performed between data from in vitro and ex vivo at 5-hours post treatment (* P < 0.05).

### Known *hipA* high-persistence mutations are absence in NRG857c

The HipA7 variant contains two mutations, G22S and D291A, associated with a high-persistence phenotype [22]. NRG857c HipA (WP_001125432) and its identical LF82 homologue both lack the key mutations of HipA7 (S4 Fig). In addition, *hipA* (D88N) and *hipA* (P86L), isolated from patients or laboratory screens, also lead to a high-persistence phenotype [23], however, none of them are present in HipA proteins from AIEC. Sequencing of the *hipA* allele from surviving colonies after 3-hour of antibiotic treatment, revealed that NRG857c did not acquired *hip*A mutations through the experiments that could explain its high-persistence phenotype (data not shown).

Although functional experiments are needed, our results show that *hipA* does not seem be necessary or sufficient to give a high-persistence phenotype to NRG857c, and other chromosomal genetic factors must be involved, as we discuss below.

## Discussion

For AIEC, the presence of persister cells had been demonstrated only for the LF82 strain [11]. In this study, we expanded the antibiotic persister analysis to other AIEC strains and revealed that this pathotype comprises a heterogeneous population with different persister cell formation competencies. One of the most significant outcomes was the identification of high-persistence strains, which differ from the LF82 behavior.

LF82 and NRG857c strains induce a distinct inflammatory response in intestinal epithelial culture cells [24] and show different colonization capabilities in mouse models [25]. Contrary to LF82, which did not colonize conventional mice for long periods, NRG857 can induce a persistent infection leading to chronic inflammation [25]. As shown in this study, it is feasible to speculate that such phenotypic differences might be due to their dissimilarities in persister cell formation. Main genetic differences between LF82 and NRG857c are localized in plasmids [18]. However, though plasmid pO83_CORR carried by NRG857c has been linked to antimicrobial peptide resistance and colonization [26], our results discard its role in the persistence phenotype (Fig 1B). Therefore, unknown actors involved in NRG857c mechanisms for persister cell formation must be chromosomal encoded and we speculate that differential gene expression or regulation of conserved genes might explain this phenotype.

Activation of stringent response through the (p)ppGpp synthase RelA and SpoT is important for *E. coli* persister cell formation [2], and LF82 persistence [11]. Unfortunately, after several unsuccessful attempts of *spoT*/*relA* deletion in NRG857c or NRG857c(Cu), we were unable to demonstrate their role on the high-persistence phenotype. However, we discarded the role of *htrA* (Fig 3C-3D) and *hipA* being responsible on this phenotype.

Recently, a *ptsI* mutation was associated to increased persister formation [27]. Our in-silico analysis revealed that NRG857c and LF82 encode identical *pstI* genes (data not shown), so gene functionality should be tested to elucidate their role in AIEC.

According to our findings, it is feasible to speculate a contribution of persister cells for NRG857c chronic colonization. Still, since genetic and phenotypic heterogenicity exists among AIEC isolates, coupled with the reduced AIEC strains analyzed here, it needs further clarification whether the high-persistence phenotype is a common characteristic among the pathotype or a particularity of specific strains.

Our study expands the known diversity among AIEC strains and underlines the importance of further studies examining the role of AIEC persisters on ongoing CD, symptoms relapse, and response to antibiotic treatment.

## Materials and Methods

### Time-killing curves

Fresh LB broth was inoculated at a starting OD_600nm_ of 0.03 from an overnight culture and growth until it reached the early exponential growth phase (OD_600nm_ 0.3-0.4). Ciprofloxacin was added to 20-80x MIC (S2 Table) to each culture and grown up to 5-hours. Samples were taken at several time points after the antibiotic challenge, serially diluted in phosphate-buffered saline (PBS, Merck), and plated on LB-agar without antibiotics. After incubation, CFU/ml were determined, and the survival ratio (regarding the number of CFU/ml at a given time to the number of CFU/ml at the treatment time) was graphed as a function of time. Time-killing curves were performed in biological triplicates.

### Macrophage-induced persisters

The J774.A1 murine macrophage cell line (ATCC TIB-67) was maintained in high glucose Dulbecco’s Modified Eagle (DMEM) medium (HyClone^TM^ Cytiva) supplemented with 10% fetal bovine serum (HyClone^TM^ Cytiva) and penicillin/streptomycin (Corning). Cells were grown at 37°C with 5% CO_2_ with regular media changes. For infection assays, macrophages were seeded at 9.5×10^5^ cells per well in 6-well plates (SPL Life Sciences) 20-24 hours prior to infection. Bacteria were grown in LB broth until the early exponential phase and then diluted in non-supplemented DMEM medium to infect macrophages at a multiplicity of infection of 10. After 10 min of centrifugation at 900xg and a 20 min incubation period at 37°C (30 min infection in total), infected macrophages were washed with PBS and lysed with 0.1% Triton X-100 (Merck). Intracellular bacteria were collected by centrifugation at 14,000xg per 2 min and resuspended in fresh LB broth, the antibiotic was added to the culture, and the time-killing curve protocol was followed as above. The survival of the macrophage-exposed population (ex vivo persisters) was compared to the survival of bacteria used as inoculum for macrophage infection (in vitro persisters).

## Authors’ contributions

PB conceptualized the study. VPV and PB designed and conducted the investigation of the study. PB wrote the original draft. VPV, RV, MAH and PB reviewed and edited the manuscript. RV and PB acquired the funding. PB supervised the study.

## Acknowledgments

We thank Alfredo Torres, Olivier Espéli, Brian Coombes and Charles Dozois for generously giving us bacterial strains and plasmids for this study. Also, we thank Laurence van Melderen and Natacha Mine for their help performing killing curve experiments.

## SUPPLEMENTAL METHODS

### Bacterial strains and growth conditions

Bacterial strains used in this study are described at S1 Table. Bacteria were grown routinely in Luria-Bertani Lennox (LB) broth (BD Difco) at 37°C with shaking at 170 rpm. When needed, 1,5 % p/v agar (LB-agar) or 15 μg/ml gentamicin (Sigma) were added.

### Bacterial growth curves in MOPS-minimal medium

Bacterial strains were inoculated from a saturated culture into MOPS-based medium supplemented with 0.4% glucose prepared as described in [1]. Growth curves were done in 96-well plates incubated in a TECAN spectrophotometer (Infinity 2000, TECAN) and started at OD_600nm_ 0.03 (to resemble killing curves conditions). Plates were incubated at 37°C with shaking, and density was measured each 10 min for a total period of 24-hours. Data from triplicates cultures were analyzed in GraphPad Prism 10 Version 10.3.0.

### Time-killing curves in MOPS-based medium

*E. coli* NRG857c overnight cultures were inoculated from frozen glycerol stocks into 2 mL of MOPS-based medium and grown overnight (18–20 hours) at 37°C with shaking at 170 rpm. Fresh MOPS-based medium was inoculated at a starting OD_600nm_ of 0.03, and growth until reach early exponential growth phase (OD_600nm_ 0.3-0.4) with good aerations. Ciprofloxacin was added to 0.15, 0.30 or 0.60 µg/ml to each culture and grown up to 5-hours at 37°C with aeration. Samples were taken at several time points after antibiotic treatment, serially diluted in PBS and plated on LB-agar without antibiotic. After incubation at 37°C, CFU/ml were determined, and the survival ratio was graphed as a function of time.

### Polymerase chain reaction (PCR)

PCR reactions were done using the SapphireAmp Fast PCR Master Mix (Takara), 0.2 μM of oligonucleotides (S2 Table) and 20 ng of pGP-Tn7-Gm or colony lysates as template. The cycling program consisted of an initial denaturation at 94°C per 1 min, 30 cycles of 98°C per 5 sec, 55°C per 5 sec, and 72°C per 20 sec, followed by a final extension at 72°C per 2 min.

### *htrA* deletion mutant construction

The AIEC strain NRG857c Δ*htrA* mutant was generated via Lambda-Red recombination [2] using the pKD46_Km recombinase-expressing plasmid [3]. Oligonucleotides koHtrA-40_G1.2 and koHtrA-40_G2.2 were used to amplify the gentamicin resistance cassette using pGP-Tn7-Gm [4] as a template. Transformants carrying pKD46_Km were transformed with the PCR product and spread onto LB-agar supplemented with gentamicin. Colonies were screened by colony PCR using oligonucleotides Up_HtrA-F and Down_HtrA-R and gene disruption was confirmed by Sanger sequencing.

### Minimum Inhibitory Concentrations (MIC)

Susceptibilities to ciprofloxacin (Sigma) were determined by broth microdilution method in Mueller-Hinton broth (BD Difco) with inocula of 5×10^5^ CFU/ml, according to CLSI M07-A10 guidelines [5]. Microplates were incubated static overnight at 37°C and MIC values were determined as the lower antibiotic concentration that inhibited growth. All MIC values were calculated from three independent experiments, involving three replicates each.

### Bioinformatic analysis

Nucleotides and protein sequences were obtained from the NCBI database. *E. coli* MG1655 HipA protein sequence (NP_416024.1) was used as a query to search by tblastn at NCBI. Aminoacid sequence alignments were generated by Clustal O (1.2.4).

### Statistical analysis

Statistical differences were determined using a two-tailed Student *t*-Test on the means of at least three independent experiments, using GraphPad Prism 10 Version 10.3.0. Differences were considered statistically significant when P < 0.05.

## SUPPLEMENTAL TABLES

**S1 Table.** Bacterial strains and plasmids used in this work.

| Strain/plasmid | Characteristics | Source |
| --- | --- | --- |
| LF82 | AIEC reference strain | gift from Olivier Espéli |
| LF82 $\Delta htrA::Km$ | LF82 *htrA* deletion mutant | gift from Olivier Espéli |
| NRG857c | AIEC reference strain | gift from Alfredo Torres |
| NRG857c $\Delta htrA::Gm$ | NRG857c *htrA* deletion mutant | This work |
| NRG857c(Cu) | NRG857c strain cured of its resistance plasmid [6] | gift from Alfredo Torres |
| H10407 | Enterotoxigenic *E. coli* (ETEC) reference strain | lab collection |
| E2348/69 | Enteropathogenic *E. coli* (EPEC) reference strain | lab collection |
| EI-34 | Enteroinvasive *E. coli* (EIEC) reference strain | lab collection |
| F-1845 | Diffusely adherent *E. coli* (DAEC) reference strain | lab collection |
| CFT073 | Uropathogenic *E. coli* (UPEC) reference strain | lab collection |
| CD1a | Clinical isolated with an AIEC phenotype [7] | lab collection |
| CD2a | Clinical isolated with an AIEC phenotype [7] | lab collection |
| CD6b | Clinical isolated with an AIEC phenotype [7] | lab collection |
| CD6r | Clinical isolated with an AIEC phenotype [7] | lab collection |
| pKD46_Km | Recombinase-expressing plasmid [3] | gift from Brian Coombes |
| pGP-Tn7-Gm | Gentamicin resistance cassette template plasmid [4] | gift from Charles Dozois |

**S2 Table.** Ciprofloxacin MIC values of bacterial strains used in this study. MIC represent mean values and SD of three independent experiments. In this study 0.6 µg/ml or 6 µg/ml ciprofloxacin (*) were used and the corresponding MIC fold for each strain is indicated at the last column.

| Strain | MIC (µg/ml) ± SD | MIC fold<br>(0.6 µg/ml)<br>(6 µg/ml)* |
| --- | --- | --- |
| LF82 | 0.0210 ± 0.0080 | 28.57 |
| LF82 $\Delta htrA::Km$ | 0.0234 ± 0.0191 | 25.64 |
| NRG857c | 0.0117 ± 0.0043 | 51.28 |
| NRG857c $\Delta htrA::Gm$ | 0.0104 ± 0.0040 | 57.69 |
| NRG857c(Cu) | 0.0160 ± 0.0000 | 37.50 |
| CD1a | 0.0078 ± 0.0000 | 76.92 |
| CD2a | 0.0078 ± 0.0000 | 76.92 |
| CD6b | 0.2188 ± 0.0579 | 27.42* |
| CD6r | 0.1250 ± 0.0000 | 48.00* |
| E2348/69 (EPEC) | 0.2500 ± 0.0000 | 24.00* |
| H10407 (ETEC) | 0.0078 ± 0.0000 | 76.92 |
| CFT073 (UPEC) | 0.0130 ± 0.0040 | 46.15 |
| EI-34 (EIEC) | 0.0078 ± 0.0000 | 76.92 |
| F-1845 (DAEC) | 0.0143 ± 0.0032 | 41.96 |

**S3 Table.**
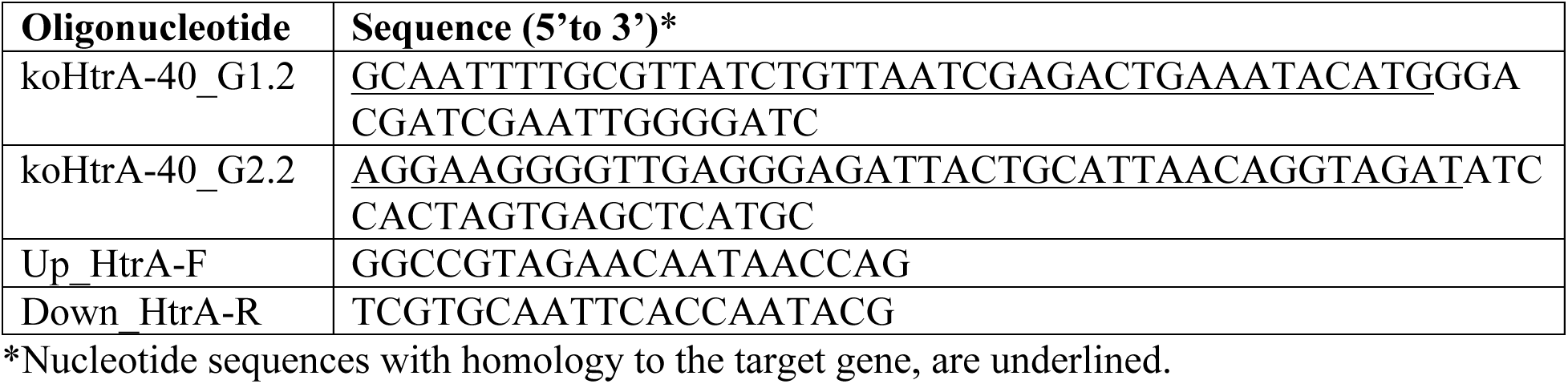
Oligonucleotides used in this study.

## SUPPLEMENTAL FIGURES

**S1 Figure.**
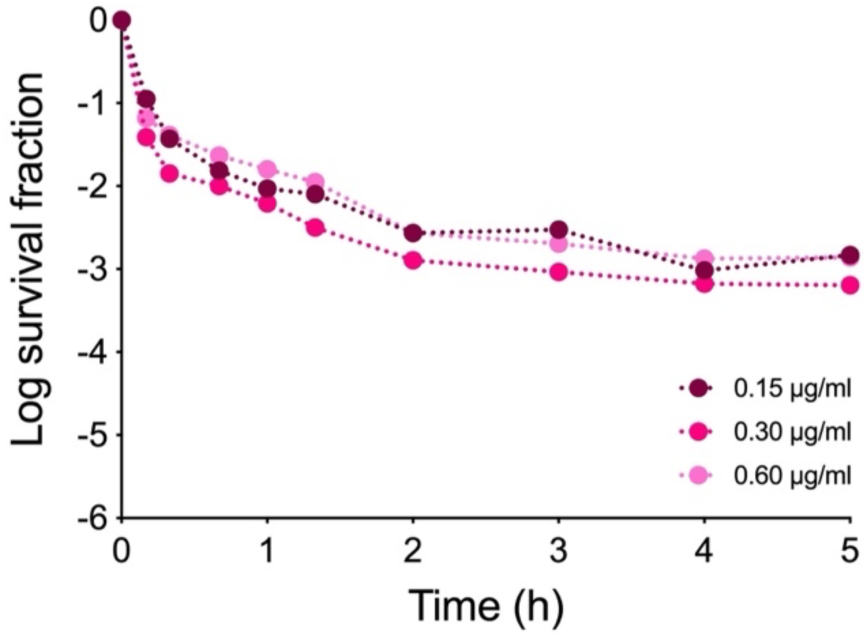
Time-killing curve of NRG857c in MOPS-minimal medium. The NRG857c strain was grown in a MOPS-based medium supplemented with 0.4% glucose until OD_600nm_ 0.3, then challenged with ciprofloxacin at 0.15, 0.30, and 0.60 μg/ml, and survival was monitored at indicated times.

**S2 Figure.**
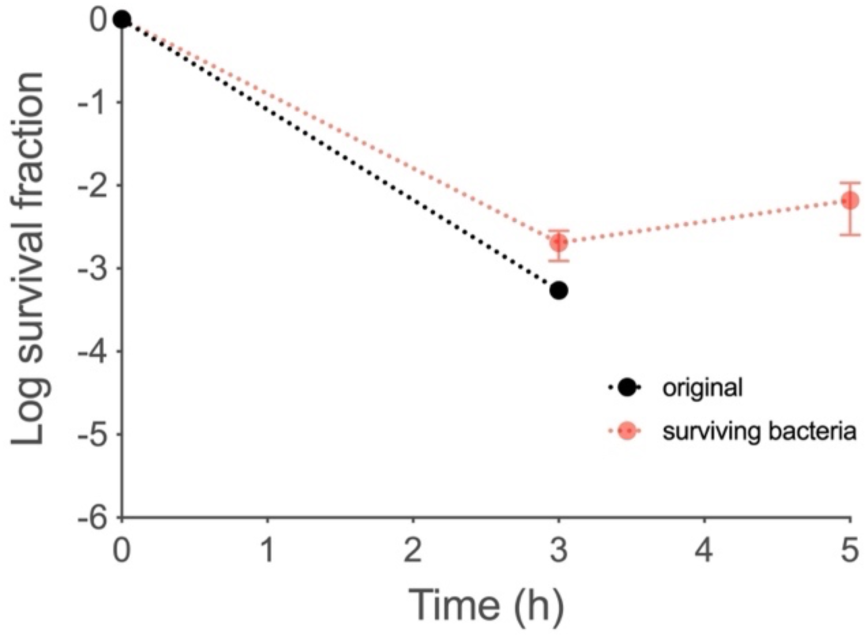
Time-killing curve of NRG857c surviving bacteria. Survival bacteria recovered after 3-hours post-ciprofloxacin challenge (50-fold MIC; black line), were grown and treated again with antibiotic at the same original conditions, and survival fraction was calculated at different time points (orange line). Data points are the mean values of three independent experiments.

**S3 Figure.**
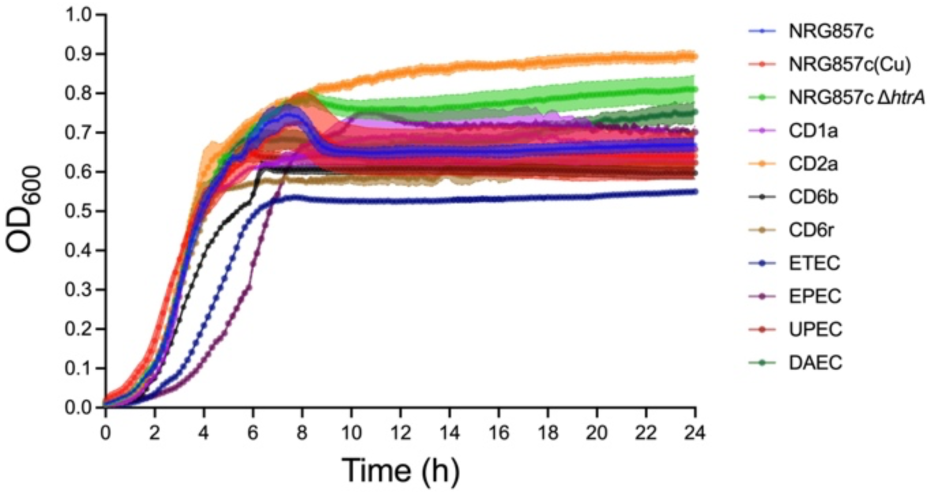
MOPS-minimal medium growth curves of *E. coli* strains used in this study. LF82, LF82 Δ*htrA,* and EIEC strains were not included, as they were unable to grow in this minimal medium.

**S4 Figure.**
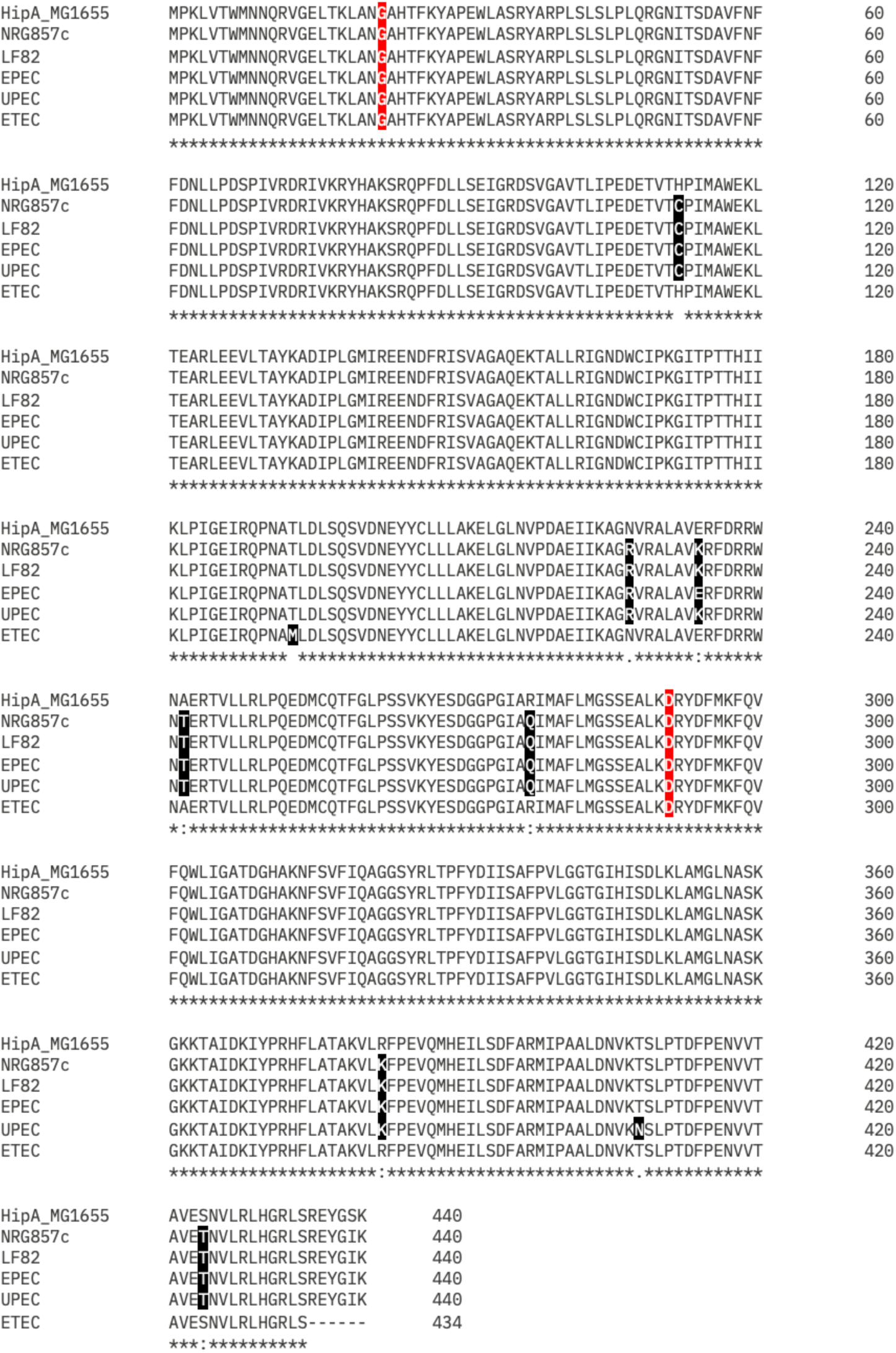
Aminoacid alignment of HipA sequences from reference *E. coli* pathotypes. Residues that differ from the canonical HipA from *E. coli* MG1655 are highlighted in black. G22 and D291 residues, which show variants in HipA7, are highlighted in red. HipA sequences accession numbers: NP_416024.1 (MG1655), WP_001125432 (NRG857c), WP_001125432 (LF82), CAS09186 (EPEC E2348/69), WP_001125431 (UPEC CFT073) and WP_001125437 (ETEC H10407).

## References

1. Moldoveanu AL, Rycroft JA, Helaine S. Impact of bacterial persisters on their host. Current Opinion in Microbiology. 2021;59: 65–71. doi:10.1016/j.mib.2020.07.006

2. Gollan B, Grabe G, Michaux C, Helaine S. Bacterial Persisters and Infection: Past, Present, and Progressing. Annu Rev Microbiol. 2019;73: 359–385. doi:10.1146/annurev-micro-020518-115650

3. Fisher RA, Gollan B, Helaine S. Persistent bacterial infections and persister cells. Nat Rev Microbiol. 2017;15: 453–464. doi:10.1038/nrmicro.2017.42

4. Helaine S, Cheverton AM, Watson KG, Faure LM, Matthews SA, Holden DW. Internalization of *Salmonella* by Macrophages Induces Formation of Nonreplicating Persisters. Science. 2014;343: 204–208. doi:10.1126/science.1244705

5. Niu H, Gu J, Zhang Y. Bacterial persisters: molecular mechanisms and therapeutic development. Sig Transduct Target Ther. 2024;9: 174. doi:10.1038/s41392-024-01866-5

6. Kim J-S, Wood TK. Persistent Persister Misperceptions. Front Microbiol. 2016;07. doi:10.3389/fmicb.2016.02134

7. Harms A, Fino C, Sørensen MA, Semsey S, Gerdes K. Prophages and Growth Dynamics Confound Experimental Results with Antibiotic-Tolerant Persister Cells. Vogel J, editor. mBio. 2017;8: e01964–17. doi:10.1128/mBio.01964-17

8. Moyed HS, Bertrand KP. hipA, a newly recognized gene of Escherichia coli K-12 that affects frequency of persistence after inhibition of murein synthesis. J Bacteriol. 1983;155: 768–775. doi:10.1128/jb.155.2.768-775.1983

9. Goneau LW, Yeoh NS, MacDonald KW, Cadieux PA, Burton JP, Razvi H, et al. Selective Target Inactivation Rather than Global Metabolic Dormancy Causes Antibiotic Tolerance in Uropathogens. Antimicrob Agents Chemother. 2014;58: 2089–2097. doi:10.1128/AAC.02552-13

10. Mulcahy LR, Burns JL, Lory S, Lewis K. Emergence of *Pseudomonas aeruginosa* Strains Producing High Levels of Persister Cells in Patients with Cystic Fibrosis. J Bacteriol. 2010;192: 6191–6199. doi:10.1128/JB.01651-09

11. Demarre G, Prudent V, Schenk H, Rousseau E, Bringer M-A, Barnich N, et al. The Crohn’s disease-associated Escherichia coli strain LF82 relies on SOS and stringent responses to survive, multiply and tolerate antibiotics within macrophages. Helaine S, editor. PLoS Pathog. 2019;15: e1008123. doi:10.1371/journal.ppat.1008123

12. Darfeuille-Michaud A, Boudeau J, Bulois P, Neut C, Glasser A-L, Barnich N, et al. High prevalence of adherent-invasive Escherichia coli associated with ileal mucosa in Crohn’s disease. Gastroenterology. 2004;127: 412–421. doi:10.1053/j.gastro.2004.04.061

13. Qiu P, Ishimoto T, Fu L, Zhang J, Zhang Z, Liu Y. The Gut Microbiota in Inflammatory Bowel Disease. Front Cell Infect Microbiol. 2022;12: 733992. doi:10.3389/fcimb.2022.733992

14. Alhagamhmad MH, Day AS, Lemberg DA, Leach ST. An overview of the bacterial contribution to Crohn disease pathogenesis. Journal of Medical Microbiology. 2016;65: 1049–1059. doi:10.1099/jmm.0.000331

15. Ledder O. Antibiotics in inflammatory bowel diseases: do we know what we’re doing? Transl Pediatr. 2019;8: 42–55. doi:10.21037/tp.2018.11.02

16. Boudeau J, Glasser A-L, Masseret E, Joly B, Darfeuille-Michaud A. Invasive Ability of an *Escherichia coli* Strain Isolated from the Ileal Mucosa of a Patient with Crohn’s Disease. Orndorff PE, editor. Infect Immun. 1999;67: 4499–4509. doi:10.1128/IAI.67.9.4499-4509.1999

17. Glasser A-L, Boudeau J, Barnich N, Perruchot M-H, Colombel J-F, Darfeuille-Michaud A. Adherent Invasive *Escherichia coli* Strains from Patients with Crohn’s Disease Survive and Replicate within Macrophages without Inducing Host Cell Death. Tuomanen EI, editor. Infect Immun. 2001;69: 5529–5537. doi:10.1128/IAI.69.9.5529-5537.2001

18. Nash JH, Villegas A, Kropinski AM, Aguilar-Valenzuela R, Konczy P, Mascarenhas M, et al. Genome sequence of adherent-invasive Escherichia coli and comparative genomic analysis with other E. coli pathotypes. BMC Genomics. 2010;11: 667. doi:10.1186/1471-2164-11-667

19. Bringer M-A, Barnich N, Glasser A-L, Bardot O, Darfeuille-Michaud A. HtrA Stress Protein Is Involved in Intramacrophagic Replication of Adherent and Invasive *Escherichia coli* Strain LF82 Isolated from a Patient with Crohn’s Disease. Infect Immun. 2005;73: 712–721. doi:10.1128/IAI.73.2.712-721.2005

20. Allen CA, Niesel DW, Torres AG. The effects of low-shear stress on Adherent-invasive *Escherichia coli*. Environmental Microbiology. 2008;10: 1512–1525. doi:10.1111/j.1462-2920.2008.01567.x

21. De La Fuente M, Franchi L, Araya D, Díaz-Jiménez D, Olivares M, Álvarez-Lobos M, et al. Escherichia coli isolates from inflammatory bowel diseases patients survive in macrophages and activate NLRP3 inflammasome. International Journal of Medical Microbiology. 2014;304: 384–392. doi:10.1016/j.ijmm.2014.01.002

22. Korch SB, Henderson TA, Hill TM. Characterization of the *hipA7* allele of *Escherichia coli* and evidence that high persistence is governed by (p)ppGpp synthesis. Molecular Microbiology. 2003;50: 1199–1213. doi:10.1046/j.1365-2958.2003.03779.x

23. Schumacher MA, Balani P, Min J, Chinnam NB, Hansen S, Vulić M, et al. HipBA–promoter structures reveal the basis of heritable multidrug tolerance. Nature. 2015;524: 59–64. doi:10.1038/nature14662

24. Eaves-Pyles T, Allen CA, Taormina J, Swidsinski A, Tutt CB, Eric Jezek G, et al. Escherichia coli isolated from a Crohn’s disease patient adheres, invades, and induces inflammatory responses in polarized intestinal epithelial cells. International Journal of Medical Microbiology. 2008;298: 397–409. doi:10.1016/j.ijmm.2007.05.011

25. Small C-LN, Reid-Yu SA, McPhee JB, Coombes BK. Persistent infection with Crohn’s disease-associated adherent-invasive Escherichia coli leads to chronic inflammation and intestinal fibrosis. Nat Commun. 2013;4: 1957. doi:10.1038/ncomms2957

26. McPhee JB, Small CL, Reid-Yu SA, Brannon JR, Le Moual H, Coombes BK. Host Defense Peptide Resistance Contributes to Colonization and Maximal Intestinal Pathology by Crohn’s Disease-Associated Adherent-Invasive Escherichia coli. Bäumler AJ, editor. Infect Immun. 2014;82: 3383–3393. doi:10.1128/IAI.01888-14

27. Parsons JB, Sidders AE, Velez AZ, Hanson BM, Angeles-Solano M, Ruffin F, et al. In-patient evolution of a high-persister *Escherichia coli* strain with reduced in vivo antibiotic susceptibility. Proc Natl Acad Sci USA. 2024;121: e2314514121. doi:10.1073/pnas.2314514121

## Supplemental References

1. Goormaghtigh F, Van Melderen L. Optimized Method for Measuring Persistence in Escherichia coli with Improved Reproducibility. In: Michiels J, Fauvart M, editors. Bacterial Persistence. New York, NY: Springer New York; 2016. pp. 43–52. doi:10.1007/978-1-4939-2854-5_4

2. Datsenko KA, Wanner BL. One-step inactivation of chromosomal genes in *Escherichia coli* K-12 using PCR products. Proc Natl Acad Sci USA. 2000;97: 6640–6645. doi:10.1073/pnas.120163297

3. Elhenawy W, Tsai CN, Coombes BK. Host-Specific Adaptive Diversification of Crohn’s Disease-Associated Adherent-Invasive Escherichia coli. Cell Host & Microbe. 2019;25: 301–312.e5. doi:10.1016/j.chom.2018.12.010

4. Crépin S, Houle S, Charbonneau M-È, Mourez M, Harel J, Dozois CM. Decreased Expression of Type 1 Fimbriae by a *pst* Mutant of Uropathogenic Escherichia coli Reduces Urinary Tract Infection. Camilli A, editor. Infect Immun. 2012;80: 2802–2815. doi:10.1128/IAI.00162-12

5. Cockerill F. Methods for dilution antimicrobial susceptibility tests for bacteria that grow aerobically: approved standard. Tenth edition. Wayne, Pa.: Clinical and Laboratory Standards Institute; 2015.

6. Allen CA, Niesel DW, Torres AG. The effects of low-shear stress on Adherent-invasive *Escherichia coli*. Environmental Microbiology. 2008;10: 1512–1525. doi:10.1111/j.1462-2920.2008.01567.x

7. De La Fuente M, Franchi L, Araya D, Díaz-Jiménez D, Olivares M, Álvarez-Lobos M, et al. Escherichia coli isolates from inflammatory bowel diseases patients survive in macrophages and activate NLRP3 inflammasome. International Journal of Medical Microbiology. 2014;304: 384–392. doi:10.1016/j.ijmm.2014.01.002

